# Disentangling objects’ contextual associations from perceptual and conceptual attributes using time-resolved neural decoding

**DOI:** 10.1101/2025.05.29.656895

**Authors:** Ariel H. Kim, Genevieve L. Quek, Denise Moerel, Olivia Gorton, Thomas A. Carlson

## Abstract

Humans effortlessly relate what they see to what they know, drawing on existing knowledge of objects’ perceptual, conceptual, and contextual attributes while searching for and recognising objects. While prior studies have investigated the temporal dynamics of perceptual and conceptual object properties in the neural signal, it remains unclear whether and when contextual associations are uniquely represented. In this study, we used representational similarity analysis on electroencephalography (EEG) data to explore how the brain processes the perceptual, conceptual, and contextual dimensions of object knowledge over time. Using human similarity judgments of 190 naturalistic object concepts presented as either as images or words, we constructed separate behavioural models of objects’ perceptual, conceptual, and contextual properties. We correlated these models with neural patterns from two EEG datasets, one publicly available and one newly collected, both recorded while participants passively viewed the same object stimuli. Across both datasets, we found that perceptual features dominated the early EEG response to object images, while conceptual features emerged later. Contextual associations were also reflected in neural patterns, but their explanatory power largely overlapped with that of conceptual models, suggesting limited unique representation of objects’ contextual attributes under passive viewing conditions. These results highlight the brain’s integration of perceptual and conceptual information when processing visual objects. By combining high temporal resolution EEG with behaviourally derived models, this study advances our understanding of how distinct dimensions of object knowledge are encoded in the human brain.

---

Relating what we see to what we know is an essential cognitive function for humans, enabling us to draw on prior experience to successfully detect, recognise and interact with objects in the world. Doing so requires rapid and reliable evaluation of the similarity between incoming sensory information and stored representations (Clarke & Tyler, 2015; Edelman, 1998; Kok et al., 2013; Nosofsky, 1986). Despite its ubiquity, this process of comparison is inherently complex, as objects themselves are defined by a myriad of both observable visual properties (e.g., colour, surface, texture, global shape, etc.) and more abstract, conceptual attributes (e.g., biological status, action affordances, abstract qualities) (Clarke et al., 2013; Clarke & Tyler, 2015; Grootswagers et al., 2019b; Hebart et al., 2020; Teichmann et al., 2026). As such, objects can be both similar *and* different to one another, depending on the dimension under consideration. For example, an alligator and a rabbit are at once both vastly disparate – one is large, scaly, and dangerous; the other small, fluffy, and cute – and yet fundamentally similar – both animals that breathe, eat, and reproduce.

Recent largescale efforts to characterise the multidimensional nature of object representations have uncovered a host of dimensions that underlie human judgements of similarity both across (Hebart et al., 2019, 2020; Teichmann et al., 2026) and within overarching object categories (Almeida et al., 2023). Findings suggest that both perceptual and conceptual attributes serve to structure object representations in the human visual system. Identified perceptual dimensions capture intrinsic properties of objects’ visual appearance (e.g., roundness, elongation, colour, flatness, shininess, texture), where conceptual dimensions encompass both taxonomic category (e.g., animal, vehicle, plant), as well as higher-level, semantically rich aspects of object meaning (e.g., associations of value, heat, disgust). Disentangling how perceptual and conceptual object properties emerge in time-resolved M/EEG neural responses has been a focus within existing work on the temporal unfolding of visual object processing (Bankson et al., 2018; Carlson et al., 2013; Clarke & Tyler, 2015; Contini et al., 2017; Giari et al., 2020; Grootswagers et al., 2019b; Isik et al., 2014; Wardle et al., 2016). This is important, since perceptual and conceptual object attributes often covary: Think of a *HAMMER* and a *FORK*, which are similar in terms of not only material composition, size, and shape, but also function, affordance and taxonomic status (graspable tools). As a collective, these studies suggest that perceptual features give rise to early, stimulus-locked activity patterns – reflecting rapid encoding of objects’ visual features – where representations of conceptual properties tend to emerge comparatively later in the neural time course, suggesting a temporal cascade from early perceptual analysis to later semantic integration (Bankson et al., 2018; Carlson et al., 2013; Clarke & Tyler, 2015; Grootswagers et al., 2019b; Proklova et al., 2019; Teichmann et al., 2026). Notably, neural representations of conceptual features also show greater variability across individuals, potentially reflecting differences in experience, familiarity, or conceptual organisation (Teichmann et al., 2026).

In addition to their inherent perceptual and conceptual features, object knowledge also comprises contextual attributes, defined by objects’ predictable associations with certain environments (e.g., a toothbrush in a bathroom, a knife on a countertop). A wealth of prior work has shown that contextual associations between objects and surrounding scenes influence how we search for and recognise objects (Bar, 2004; Biederman et al., 1973; Davenport & Potter, 2004; Ganis & Kutas, 2003; Oliva & Torralba, 2007; Võ et al., 2019), with scene context even serving to disambiguate neural representations for degraded objects that are otherwise unrecognisable (Brandman & Peelen, 2017; Wischnewski & Peelen, 2021). Context-based facilitation also arises at the inter-object level, with observers recognising objects more efficiently when they are contextualised by second, related object (Auckland et al., 2007; Bar & Ullman, 1996; Davenport, 2007; Green & Hummel, 2006; Kaiser et al., 2019; Kaiser et al., 2014; Quek et al., 2025). Moreover, violations of canonical contextual associations seem to elicit reliable neural signals even when observers are not explicitly attending to congruity between the elements (Mudrik et al., 2010, 2014; Quek & Peelen, 2020), suggesting that representations of contextual object attributes may be evoked automatically during object processing.

Where prior work characterising the neural timecourse of individual object dimensions does encompass aspects of objects’ contextual attributes (e.g., ‘outdoors-related’, ‘bathroom-/wetness related’, Hebart et al., 2023), to date no study has explicitly aimed at *dissociating* time-resolved representations of contextual object knowledge from their perceptual and conceptual counterparts. Understanding the degree of temporal overlap in the processing of these three classes of object features is central to the interpretation of inter-object contextual facilitation effects, which are underpinned by both conceptual and contextual associations between simultaneously presented objects (e.g., *teacup* and *saucer* are both food/drink related and typically encountered in the same surrounding context). The present study addressed this gap by examining the extent to which behaviourally-derived models encompassing objects’ perceptual characteristics, conceptual attributions, and contextual associations predict unique variance in the unfolding neural response to object images reflected in EEG data.

To explore the intersection between processing of objects’ contextual, perceptual, and conceptual features, we adapted the “odd-one-out” task used by Hebart et al. (2020) to capture subjective similarity judgments among triplets of object concepts. Crucially, rather than eliciting purely intuitive judgements of object similarity, here we imposed specific task instructions designed to selectively weight a particular object dimension. Different groups of online observers made triplet similarity judgements for the same 190 object concepts in terms of *i)* objects’ observable perceptual properties (e.g., size, colour, shape, etc.), *ii)* their conceptual attributes (e.g., function, use, affordance), or *iii)* their contextual associations (e.g., where the object would likely be found). These dimension-specific judgments were then used to construct models that isolated perceptual, conceptual, or contextual structure of objects. Our overarching goal was to examine the extent to which each of these behaviourally-derived object dimension models explained unique variance in the dynamically evolving representational structure of visual object processing. To this end, we correlated our object dimension models with two high-temporal resolution EEG datasets recorded in separate participant samples viewing the same object images under orthogonal task conditions. These included both a publicly available EEG dataset (referred to as Experiment 1; (Grootswagers et al., 2022) and a newly acquired dataset (referred to as Experiment 2) which differed in presentation rate and the number of object image repetitions.

Secondarily, we also aimed to investigate how different presentation modalities would differentially capture subjective similarity between objects’ perceptual, conceptual, and contextual dimensions. Thus, we derived object dimension models based on both image and word-label versions of our 190 object concepts. This represents a departure from previous studies that have typically modelled object dimensions based on behavioural responses to object images alone (e.g., Cichy et al., 2019; Giari et al., 2020). Here we predicted that similarity judgments using word labels would capture object dimensions in a more abstract form, whereas judgments based on images would inherently be more visually-driven, even if participants were asked specifically to consider conceptual or contextual object properties. By using both abstractly derived (word-based models) and perceptually derived (image-based models) object dimensions to understand neural responses to objects, we aimed to further delineate the contributions of perceptual and conceptual dimensions to neural responses. Our analysis measured the degree to which the representational structure of object dimensions, as perceived by individuals, aligned with the dynamically evolving representational structure of visual object processing in the brain.

## 2. MATERIALS AND METHOD

The current study consisted of three parts: (1) collection of behavioural data, (2) analysis of both pre-collected EEG data (Experiment 1) and newly acquired EEG data (Experiment 2), and (3) Representational Similarity Analysis using data from (1) and (2).

### 2.1 Stimuli

To investigate how different dimensions of human similarity judgments are reflected in the neural processing of visual objects, we took advantage of the THINGS stimulus database (Hebart et al., 2023; Hebart et al., 2019). This open-source database consists of 1,854 object concepts systematically sampled from a wide range of concrete nouns that can be visually depicted and labelled in American English. These object concepts correspond to basic-level categories such as *dog, car, spaghetti,* and *foot*. Each concept is associated with a word label and includes exemplar images that showcase the objects embedded within naturalistic scenes.

These exemplar images capture the objects in diverse colours, shapes, angles, positions, lighting conditions, and other visual characteristics, creating a rich and varied visual context for each object concept. Additionally, each concept belonged to one of the 27 high-level categories defined by THINGS, which represent broader superordinate categories such as vehicles, animals, food, and body parts (Figure 1A).

**Figure 1.**
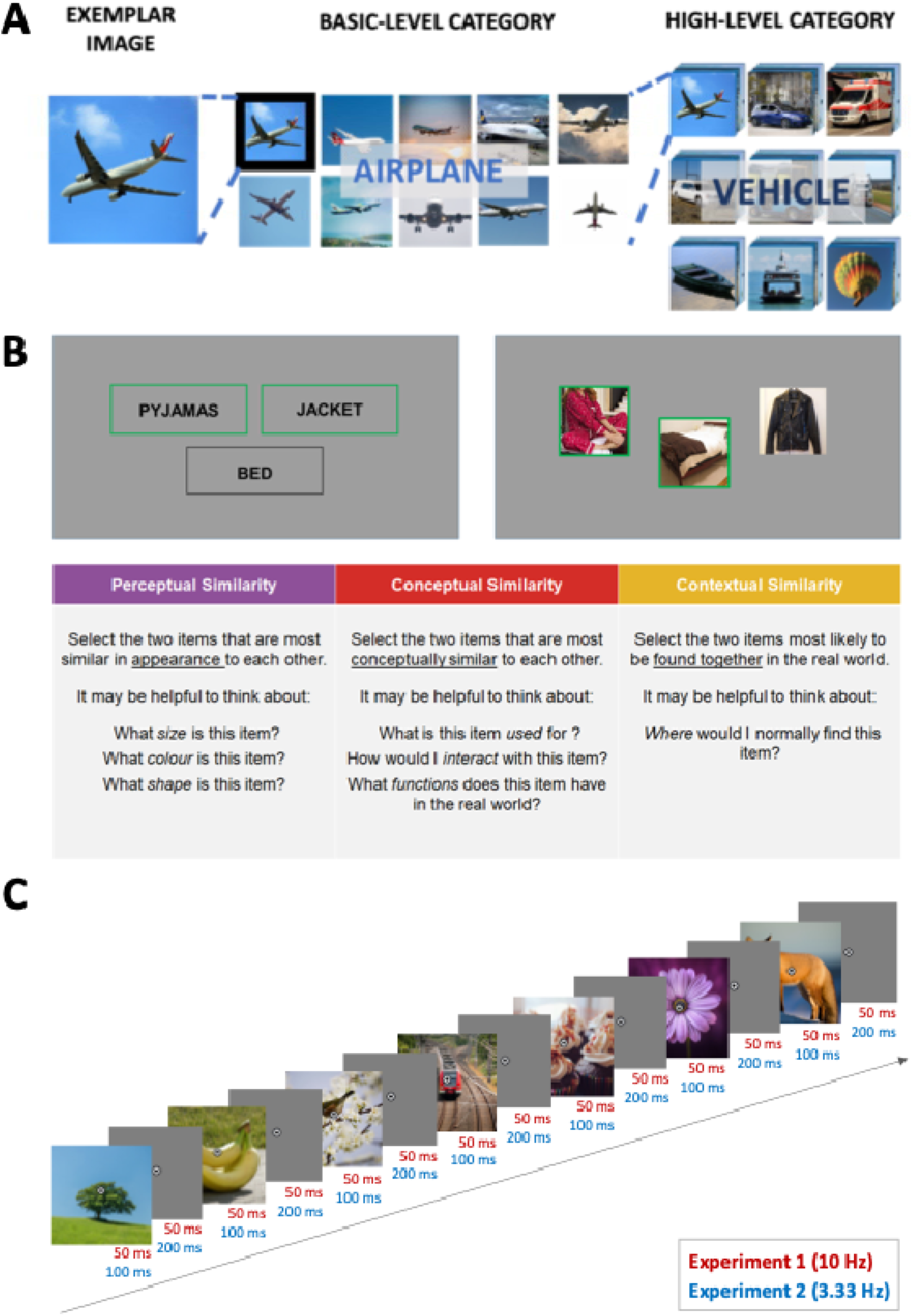
**A.** Stimuli were 190 object concepts from the THINGS database (Hebart et al., 2019). Each was represented by 10 exemplar images; there were 7-8 object concepts from each of the top 25 high-level categories identified in the original Hebart paper (2019). **B.** Overview of the triplet similarity behavioural task(s). Each participant was assigned to either a WORD or IMAGE version of the task, and instructed to evaluate triplets of object concepts along one of three objects dimension (fixed across the experiment). Thus, the participant samples for each combination of Stimulus Type and Similarity Type were non-overlapping. **C.** An example of the Rapid Serial Visual Presentation (RSVP) paradigm used in the two EEG experiments that differed in their image presentation rate (i.e., 3.33 Hz vs. 10 Hz). All images in this figure have been replaced with visually similar images from the public domain.

Since it was infeasible to derive six separate models for the full set of object concepts in the THINGS database, here we focused on a subset of 190 object concepts spanning the top 25 high-level categories identified in the original THINGS database (*food, animal, clothing, tool, sports equipment, vegetable, vehicle, musical instrument, fruit, body part, dessert, toy, container, part of car, weapon, bird, furniture, kitchen tool, office supply, kitchen appliance, plant, insect, home décor, medical equipment, electronic device*). From these high-level categories, we selected 7-8 concepts per category, which resulted in 190 object concepts in total, each of which was represented by 10 exemplar images and a word label. This resulted in a final stimulus set of 1900 images (used in behavioural and EEG data collection) and 190 words (used in behavioural data collection only).

### 2.2 Behavioural Data Collection

The goal of our behavioural data collection was to model functionally-derived similarity in our set of 190 objects’ perceptual, conceptual, and contextual attributes. To this end, we tasked different groups of observers with evaluating similarity among triplets of object concepts along one of those three dimensions. Separate models were derived for the same 190 object concepts represented by 1) images and 2) object labels (i.e., words). Similarity judgements from each task were converted into a representational dissimilarity matrix (RDM) to create six behavioural model RDMs, to be subsequently related to neural RDM series based on EEG data recorded in a different set of participants viewing image versions of same 190 object concepts.

#### 2.2.1 Participants

A total of 695 undergraduate students at the University of Sydney, Australia, were recruited to take part in the online behavioural study in exchange for course credit. Each participant took part in only one of the six behavioural tasks, and each task had a different number of participants: Perceptual-image task (n=128), Conceptual-image task (n=101), Contextual-image task (n=116), Perceptual-word task (n=113), Conceptual-word task (n=119) and Contextual-word task (n=118). All participants were aged over 18 years; no further demographic information was collected. All participants provided informed consent prior to commencing the experimental task.

#### 2.2.2 Behavioural Task Paradigm

We collected behavioural similarity judgements for our 190 object concepts using an adapted version of the “triplet odd-one-out task” from Hebart et al. (2020), in which observers evaluate the similarity across three object concepts on each trial. In Hebart et al.’s original version of this task, observers saw three object images per trial, and simply selected the object they intuitively felt is the ‘odd-one-out’, which implies that the other two objects were more similar. In our adapted version of this task, we instructed online observers to consider triplets of object concepts with respect to either their *Perceptual*, *Conceptual,* or *Contextual* features, <u>selecting the two objects that were most similar along the specified dimension</u>. The task instructions were worded differently depending on the type of similarity judgement required for each task (see Figure 1B). In the *Perceptual* condition, the instructions were: *“Select the two items that are most similar in <u>appearance</u> to each other. It may be helpful to think about: What size is this item? What colour is this item? What shape is this item?”.* In the *Conceptual* condition, the prompt was: “*Select the two items that are most <u>conceptually similar</u> to each other. It may be helpful to think about: What is this item used for? How would I interact with this item? What functions does this item have in the real world?*”. In the *Contextual* condition, participants were shown: “*Select the two items that you would be most likely to <u>find together</u> in the real world. It may be helpful to think about: Where would I normally find this item?*”. These instructions were designed to constrain observers’ similarity judgements to the specific dimension of interest, with the understanding that the same trio of objects could be evaluated differently depending on the dimension under consideration (e.g., a hammer and a fork are similar in terms of both their perceptual AND conceptual attributes, but less so in their contextual associations).

To reinforce these instructions and thus reduce the risk of participants applying inconsistent or unintended criteria, we reiterated the object dimension the participant should (and should not) attend to across two worked demonstration trials. As one example, in the contextual condition, participants were shown *PARROT, CHICK*, and *TRACTOR*, and instructed: “Remember to choose the items that <u>go together</u> in the real world. Don’t make the mistake of choosing items that are similar concepts! The correct answer is **chick** and **tractor**, which could both be found on a farm. Don’t get distracted by **chick** and **parrot** being similar concepts (both birds)!”. After examining these demonstration trials, participants completed five practice trials that were individually tailored to each version of the task, and included explicit feedback about the selected pair of objects. For example, selecting *ELEPHANT* and *BEAR* on a perceptual similarity practice trial resulted in the feedback text: “That was an easy one! An elephant and a bear both have four legs, a tail, eyes, etc.”. Conversely, selecting *ELEPHANT* and *BAMBOO* resulted in: “Hmm I’m not so sure about that one. They don’t really share many visual features”.

Participants were assigned to either a *word* or *image* format of the task (see Figure 2A), and were assigned to only *one* of the three dimensions to judge object similarity. Thus, there were six different triplet similarity tasks with unique participant samples (Perceptual-word, Conceptual-word, Contextual-word, Perceptual-image, Conceptual-image, and Contextual-image). In the word task, three words were randomly drawn from the bank of 190 labels that corresponded to the names of 190 object concepts (i.e., three distinct object concept labels per trial). In the image task, we selected three different object concepts at random, and presented the first exemplar image for each after shuffling the set of 10 exemplars. Participants in the word version of the task completed 250 trials; participants in the image task completed 170 trials. This difference was due to a practical constraint, since images took longer to load in the online study. Alongside the experimental trials, each triplet similarity task also included 5 attention-check trials, designed to probe whether the participants were correctly attending to the instructed dimension. These trials were tailored for each version of the behavioural experiment, such that each trial had an obvious answer with respect to the instructed object dimension (e.g., a perceptual similarity attention-check trial would contain *MOTH, DRAGONFLY*, and *DIRTBIKE*). The total testing time in both versions was approximately 30 minutes.

**Figure 2.**
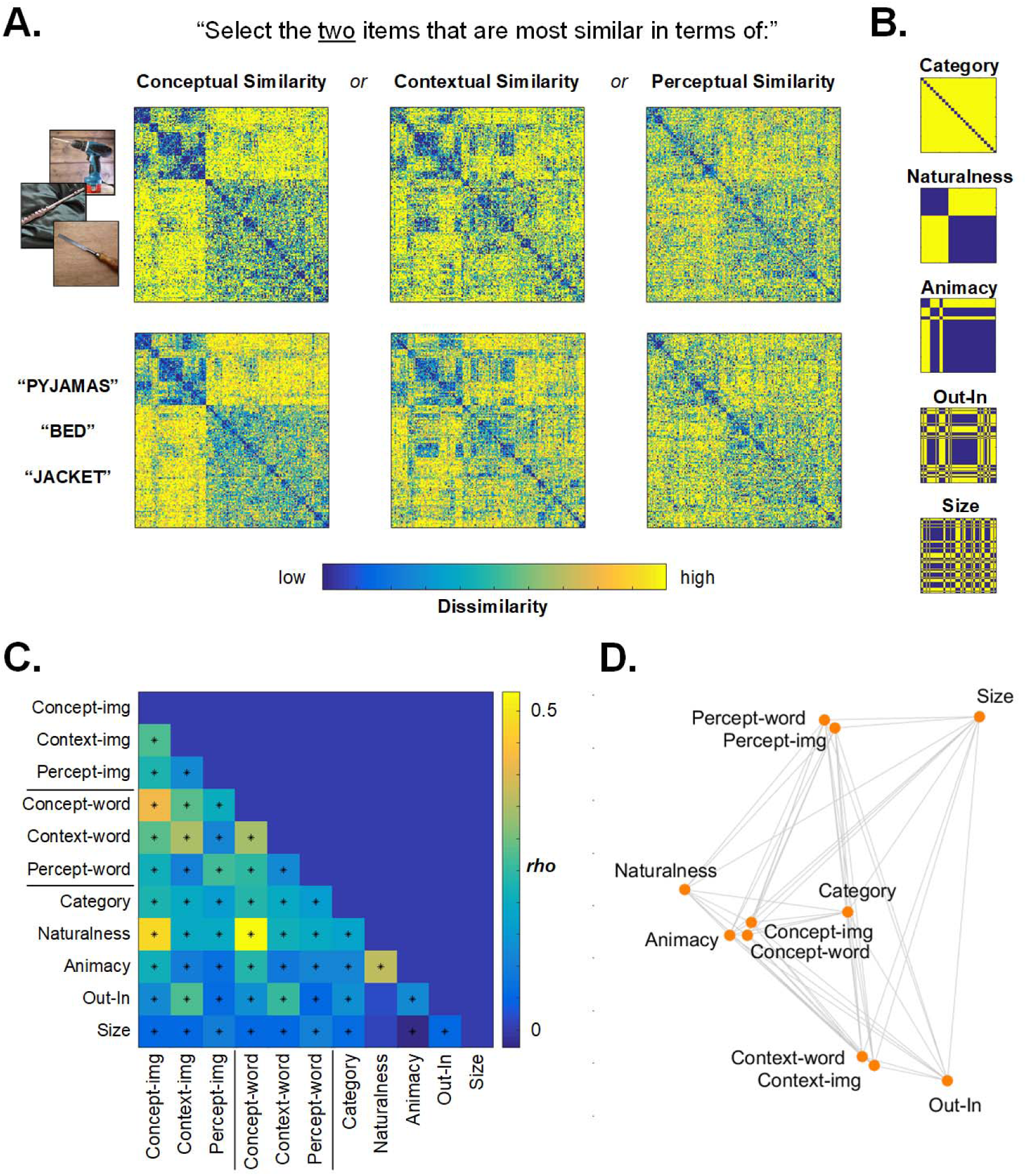
A) Behavioural RDMs derived from each of the 6 behavioural triplet similarity tasks. Non-overlapping groups of online observers saw either triplets of object-images or object-names and had to select the pair with the highest conceptual similarity, contextual similarity, or perceptual similarity. B) Binary models for the 190 object concepts. C) Intercorrelation matrix for the six behavioural RDMs and the binary models. Stars indicate correlations significant at p < .001, FDR corrected. D) Model similarity reflected in 2D multidimensional scaling. Note the close proximity of word/image versions for each object similarity dimension.

Note that for all six triplet similarity tasks, each participant was tested with only a subset of the possible triplet combinations from the 190 objects, as testing all 112,835 possible combinations per participant was practically unfeasible. The large number of possible combinations also meant that, even across all participants, we were unable to obtain results from all triplet combinations. However, previous studies have shown that testing a subset of triplet combinations still provides sufficient coverage to infer most of the pairwise dissimilarities between objects, which is necessary for constructing the model RDM (e.g., Hebart et al., 2020; Moerel et al., 2024). The number of trials collected from each task were as follows: Perceptual-image task = 21,760, Conceptual-image task = 17,170, Contextual-image task = 19,720, Perceptual-word task = 28,250, Conceptual-word task = 29,750 and Contextual-word task = 29,500. Although these represented only a fraction of the total possible trials, each trial provides dissimilarity information about three objects (i.e., A-B, A-C, and B-C), allowing for three pairwise comparisons within a single trial. Thus, a relatively small amount of trials was still able to cover a large number of unique object pairs. All behavioural data and relevant analysis code can be accessed at: https://osf.io/jy284/.

### 2.3 EEG Data Collection

The current study employed two EEG datasets recorded in response to the same set of 1900 object images, covering 190 concepts. In <u>Experiment 1</u>, we took advantage of a pre-existing, publicly available dataset from Grootswagers et al. (2022), comprising 64 channel EEG recordings obtained in 50 observers as they viewed a rapid stream of object images presented at a rate of 10 Hz (image duration = 50 ms; ISI = 50 ms; 1 repetition per image, see Figure 1C). Acquisition details for this Experiment 1 neural dataset are provided in Grootswagers et al. (2022). In <u>Experiment 2</u>, we acquired a novel dataset for this study, comprising 128 channel EEG recordings obtained in 20 observers as they viewed object images appearing at a rate of 3.33 Hz (image duration = 100 ms; ISI = 200 ms; 3 repetitions per image, see Figure 1C). The sections below provide detailed acquisition information pertaining to this Experiment 2 dataset (example dataset available at: https://osf.io/jy284/).

#### 2.3.1 Participants

Twenty undergraduate students (15 females and one non-binary individual) from the University of Sydney participated in Experiment 2 in exchange for course credit (mean age = 21.85 years, SD = 4.55, age range = 18-34). All participants reported normal or corrected-to-normal vision and normal colour vision, and had no history of psychological or neurological disorders. All but one reported being right-handed. Verbal and written informed consent was obtained from all participants before the experiment, and the study was approved by the Human Ethics Committee of the University of Sydney (ethics ID 2019/340). One participant was excluded from the results due to a technical error in the EEG data, resulting in a final sample of 19 participants.

#### 2.3.2 Experimental Design and Procedure

Experiment 2 was modelled on the paradigm employed by Grootswagers et al. (2022) to collect the EEG dataset comprising Experiment 1. Where their paper presented multiple exemplars of all 1,854 object concepts from the THINGS dataset (Hebart et al., 2019), here participants viewed a rapid stream of object images representing our selected subset of 190 concepts used in the behavioural experiment. We modified the image presentation rate used by Grootswagers et al. (2022) from 10 Hz to 3.33 Hz (see Figure 1C), on the rationale that a slower presentation rate might enable better capture of not only visual and perceptual processing of objects but also activation of more abstract and semantic information (Carlson et al., 2013; Contini et al., 2017). We selected the first 10 unique images for each per concept, and each of the 1900 object images appeared three times of the course of the full experiment.

Participants sat in a dimly lit room at a distance of ∼60 cm before a 1920 x 1080 pixel computer monitor. After being set up for EEG recording, they rested their hands on a button box and were instructed to maintain their gaze on a small, centrally-displayed black and white bullseye while images were flashing in the background. Participants were instructed to remain still and minimise eye and head movements during the stimulus presentation. Stimuli were presented on a uniform background using the PsychoPy toolbox in Python (Peirce et al., 2019), and were 256 by 265 pixels (approximately 6.25 by 6.25 degrees of visual angle) in size. The task of the participants was to press the button whenever the bullseye turned red (colour changes lasted 100 ms and occurred at random intervals 2-4 times per block). The purpose of this task was to increase attention and engagement. Before the start of the experiment, a practice trial was conducted to ensure that participants understood the task requirements.

All 190 object concepts were presented in a randomised order within each individual sequence; at a rate of 3.33 Hz, this meant that a single sequence lasted 57 seconds. There were 10 exemplar images for each object concept, with each exemplar presented three times across the entire experiment (i.e., 30 sequences in total). This resulted in a total of 5,700 stimulus presentations per participant (190 concepts x 10 exemplars x 3 repetitions). Participants were encouraged to take sufficient breaks between sequences to mitigate screen fatigue. The progress of the experiment was indicated on the screen between sequences by displaying the number of the upcoming sequence (e.g., "2/30", "18/30"). The entire experiment, including breaks, took approximately 35 minutes.

#### 2.3.3 EEG Recording and Preprocessing

During the stimulus presentation, participants’ brain activity was continuously recorded using a BrainVision ActiChamp system at a sampling rate of 1000 Hz. The standard 128-channel EEG cap (Brain Products; GmbH, Herrsching, Germany) was fitted to the participants, corresponding to the 10-20 international system of electrode locations (Oostenveld & Praamstra, 2001). We applied electrolyte gel to keep the impedances below 10 kΩ, and the electrodes were referenced to FCz.

The offline preprocessing of EEG data was performed using the EEGLAB toolbox in MATLAB (Delorme & Makeig, 2004). A Hamming windowed FIR filter was applied with a low-pass filter of 100 Hz and a high-pass filter of 0.1 Hz. The data were then re-referenced to the average reference and downsampled to a rate of 250 Hz. Epochs were created for each stimulus presentation from -100 ms to 800 ms relative to the stimulus onset. No further preprocessing steps were taken.

### 2.4 Representational Similarity Analysis

To assess whether the behavioural data of object similarity judgements could explain the dynamics of neural representational structure over time, we used Representational Similarity Analysis (RSA; Kriegeskorte et al., 2008). We created Representational Dissimilarity Matrices (RDMs), which capture the dissimilarity between pairs of experimental stimuli based on their representational patterns. For the behavioural data, RDMs were generated from participants’ similarity judgments of objects, with one RDM created for each of six triplet similarity tasks. As for the neural data, RDMs reflected the similarity or dissimilarity in the neural activation patterns evoked by each pair of the 190 concepts.

#### 2.4.1 Behavioural model RDMs

Before creating the behavioural model RDMs, we firstly removed any participants who made an incorrect selection on more than 3 of the 5 attention-check trials, on the rationale that they were not correctly attending to the instructed object dimension. This resulted in the following final participant samples: Perceptual-image task (n = 114), Conceptual-image task (n = 100), Contextual-image task (n = 108), Perceptual-word task (n = 98), Conceptual-word task (n = 105) and Contextual-word task (n = 103). Following this, we constructed 6 RDMs based on the six different triplet tasks. These RDMs were constructed at the level of object concepts, with cells reflecting the pairwise dissimilarity between objects’ selection rates (ranging between 0 and 1). This value was computed from the triplet similarity task data, by calculating the proportion in which two objects were chosen together out of all the instances when they were shown together in a triplet. A value of 1 indicated extreme dissimilarity, meaning that the two objects were never chosen together, while a value of 0 indicated extreme similarity, meaning the objects were always chosen together. As outlined above, because we presented only a subset of the total possible triplet combinations in our behavioural tasks, some cells of these RDMs had no dissimilarity score, indicating that the respective object pairs were never shown together in a triplet. The proportion of missing dissimilarity values in the six behavioural model RDMs were as follows: Perceptual-word = 1.65%, Conceptual-word = 0.52%, Contextual-word = 0.60%, Perceptual-image = 1.69%, Conceptual-image = 7.89%, and Contextual-image = 6.22%.

We inspected the validity of our behaviourally-derived models by correlating them with binary models representing ground truth object properties. These included models of high-level category, naturalness, animacy, outdoor/indoor association and real-world size (smaller or larger than a shoebox). Each object pair was given a binary value of either 0 or 1, indicating identicality or difference in the given object dimension (Figure 2B). As is clear in Figure 2C, there were significant correlations among most of our models, with the strongest correlations emerging between the Conceptual-word and Naturalness models (*r* = .53, *p*<.00001), and the Conceptual-image and Naturalness models (*r* = .48, *p* < .00001). Notably, strong correlations were evident between the image and word versions of each similarity judgement task, reflected in the clustering of these models when visualised using 2D multidimensional scaling (Figure 2D).

#### 2.4.2 Experiment 2 Neural RDMs

We subjected the preprocessed EEG data from Experiment 2 to RSA to produce an RDM series reflecting time-resolved pairwise dissimilarity between all 190 object-concepts. Decoding accuracy served to index the degree of dissimilarity between object concepts’ neural activation patterns, with higher decoding accuracy taken to reflect more dissociable object representations. In line with previous studies (e.g., Grootswagers et al., 2019), we ran the decoding analysis on preprocessed EEG data collected from all 128 channel voltages, using the CoSMoMVPA MATLAB toolbox (Oosterhof et al., 2016). A linear discriminant analysis (LDA) classifier was used to identify the decision boundary that could best differentiate the brain activation patterns evoked by objects. We implemented a leave-one-sequence-out cross-validation procedure, wherein the classifier was trained iteratively on all sequences of EEG data except one, which was kept aside to serve as the test data. We repeated this process for all sequences to ensure that every sequence was used for testing. During training, the classifier was tasked with distinguishing the neural activity patterns evoked by pairs of object objects and classifying them into two categories. This process was performed for every possible pairwise combination of the 190 object concepts shown in that sequence. The classifier underwent training and testing at each timepoint in the epoch between -100 and 800 ms. This process was repeated for each participant, resulting in one time-varying neural RDM per participant for each time-point.

#### 2.4.3 Experiment 1 Neural RDMs

To create corresponding RDMs based on the pre-existing EEG dataset for Experiment 1, we applied the same pre-processing and analysis steps described above for Experiment 2 to the raw data from Grootswagers et al. (2022). This EEG dataset consisted of EEG recordings from 50 participants presented with 1,854 THINGS concepts. Four participants were excluded from the analysis due to poor signal quality or equipment failure, resulting in data from 46 participants. From this original dataset, we extracted the data pertaining to the first 10 images of the 190 object concepts, aligning with the concepts and stimuli used in our behavioural study and the collection of new EEG data in Experiment 2. We used the same epoch length described for Experiment 2 (-100 ms to 800 ms relative to stimulus onset, or 225 time-points). We then used pairwise decoding with a leave-one-sequence-out cross-validation procedure to obtain an RDM for each participant and timepoint in the epoch.

#### 2.4.5 Correlation between Behavioural Model RDMs and EEG Neural

##### RDMs

To examine the degree to which our behaviourally-derived RDMs could account for unique variance in the neural responses elicited by images of those same object concepts, we correlated each of the six behavioural RDMs with the time-varying neural RDM from Experiment 1 and Experiment 2. For this analysis, we grouped the behavioural models by whether they were derived from image-based or word-based similarity judgements. Thus, four sets of partial correlation analyses were conducted: (1) Experiment 1 neural RDM with three image model RDMs (i.e., Perceptual-image, Conceptual-image, Contextual-image), (2) Experiment 1 neural RDM with three word model RDMs (i.e., Perceptual-word, Conceptual-word, Contextual-word), (3) Experiment 2 neural RDM with three image model RDMS, and (4) Experiment 2 neural RDM with three word model RDMs.

In each case, we were interested in quantifying the unique variance in the neural RDM that a given model RDM could explain while controlling for the other models in that set. Within each analysis and at each timepoint, we used MATLAB’s *partialcorr* function to obtain Spearman’s rank correlation coefficients between the neural RDM and a given model RDM while partialling out the variance explained by the other two model RDMs within the same task modality. This procedure serves to regress out variance shared with competing models from both the model predictions and the neural data. The resultant correlation between these residuals serves to capture the model’s independent explanatory power, rather than variance attributable to shared structure between models. Notably, we computed partial correlations at the level of the RDMs (model to neural), rather than at the level of EEG signals or stimulus-wise classification accuracies. Due to the lack of behavioural data for one object (’spatula’), we excluded it and only correlated 189 objects in both the neural and model RDMs. Additionally, as some cells in the model RDMs did not have dissimilarity scores, we used a geometric reconstruction method to complete these missing dissimilarity scores (Moerel & Grootswagers, 2025) before using them in the correlation analysis. For detailed information on this analysis, see the publicly shared analysis code available here: https://osf.io/jy284/.

Finally, we calculated difference scores between the resultant partial correlation values associated with each model using a simple arithmetic subtraction. For example, difference scores were calculated between the partial correlations of the perceptual and conceptual models within the same task (i.e., Perceptual-image minus Conceptual-image; Perceptual-word minus Conceptual-word), and between image and word modalities within the same similarity dimension (i.e., Perceptual-image minus Perceptual-word; Conceptual-image minus Conceptual-word).

### 2.5 Statistical Inference

We evaluated evidence for positive correlations between the behavioural models and the two EEG dataset using Bayesian hypothesis testing that considered evidence supporting the alternative hypothesis (i.e., H_a_) over the null hypothesis (i.e., H_0_). In this context, we report Bayes factors (BFs) which assess the likelihood of a partial correlation (H_a_) as compared to no partial correlation (H_0_), given the set of obtained correlation values across participants at each time point. Specifically, we defined H_a_ as the existence of a positive partial correlation between the neural and model RDMs (i.e., as the dissimilarity between two stimuli increases in the neural RDM, the dissimilarity in the behavioural model RDM also tends to increase), while H_0_ posited that the partial correlation between neural and model RDMs was zero (i.e., no relationship between the dissimilarity of two stimuli in the neural RDM and the model RDM). In general, a BF greater than 1 indicates evidence in favour of H_a_, while a BF less than 1 suggests evidence in favour of H_0_. We interpreted BF thresholds as follows: BF > 10 was considered strong evidence, and BF > 3 was considered weak evidence in favour of H_a_. Conversely, BF < 1/10 indicated strong evidence, and BF < 1/3 indicated weak evidence in favour of H_0_. When 1/3 < BF < 3, this was interpreted as insufficient evidence for either hypothesis. We used a threshold of BF > 3 to calculate the onset time of above chance correlations. We required at least two consecutive BFs > 3 for the onset, an isolated BF at a single time point, meeting the threshold of 3 or 1/3 but without consecutive BFs reaching these thresholds, was treated as inconclusive evidence for either hypothesis.

BFs were calculated using the Bayes Factor package in R (Morey et al., 2015). The null hypothesis was defined using a point null (Morey & Rouder, 2011) and we applied a half-Cauchy prior for the alternative hypothesis to conduct a directional test to test for positive partial correlations. The prior range for the alternative hypothesis was defined from d = 0.5 to d = ∞ to exclude small effects between the interval of d = 0 and d = 0.5, as these have been shown to occur at chance under the null hypothesis (Teichmann et al., 2022). The prior width was set to a medium (r = 0.707), which is the default setting in the BayesFactor R package.

## 3. RESULTS

We first examined whether brain activity during passive viewing of rapid object image streams could be explained by our behavioural models of objects’ perceptual, conceptual, and contextual attributes. We separated the models by the modality they were derived from – i.e., whether similarity judgements were made for object-labels (words) or object images. Within each model grouping, we computed partial correlations between the perceptual, conceptual, and contextual models and time-resolved EEG data, with the aim of identifying timepoints where each model uniquely explained variance in the EEG signal (i.e., beyond what was captured by the other models).

### 3.1 Correlating the behavioural models with the Experiment 1 neural

#### RDM

Figure 3 shows the partial correlations between the Experiment 1 EEG dataset (acquired at a 10 Hz image presentation rate) and the three object dimension models derived from the image-based triplet similarity task (Figure 3A) and the word-based triplet similarity task (Figure 3B). We observed partial correlation between the Experiment 1 neural RDM and the Perceptual-image model beginning 108 ms after stimulus onset and peaking at 192 ms. There was also a comparatively later partial correlation with the Conceptual-image model beginning from 184 ms, which coincides with its peak at 188 ms. There was evidence for no positive partial correlation with the Contextual-image model.

**Figure 3.**
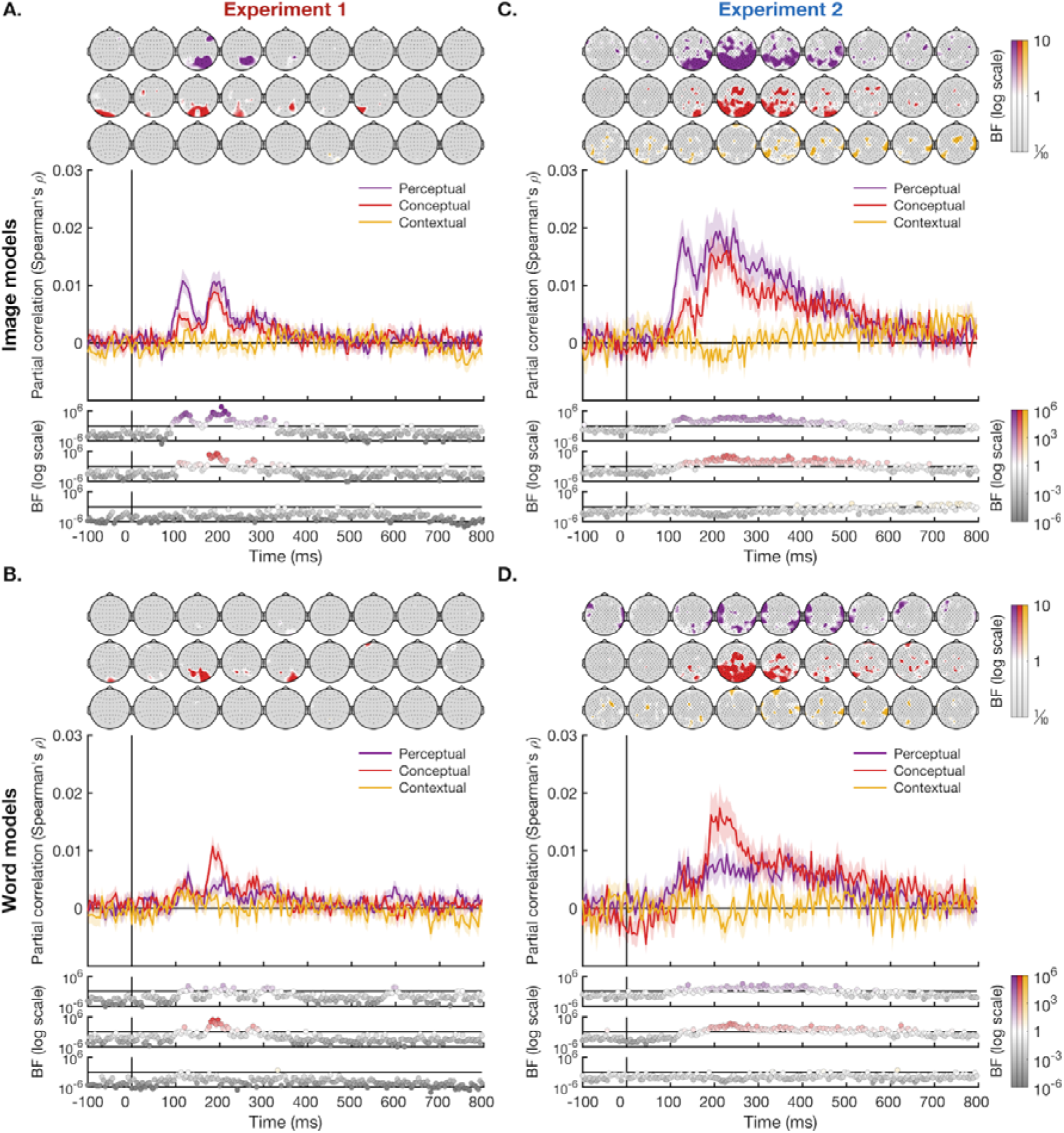
Partial Correlations Between Behavioural Models and two EEG Datasets. Partial correlations (Spearman’s *ρ*) were calculated between the EEG neural RDMs from two datasets (reflected in columns) and the six behavioural model RDMs grouped by modality (reflected in rows). Partial correlations were computed at each time point from stimulus onset by removing the variance explained by the other two models within the same modality from each model’s full correlation. Shaded areas represent standard errors. Time (ms) is relative to the stimulus onset at 0 ms. The logarithmic scale of Bayes factors (BFs) below each plot indicates the strength of evidence for two opposing hypotheses at each individual timepoint. BF>1 were represented by circles filled with the corresponding plot colour and denote evidence for alternative hypothesis (i.e., positive or negative correlation), BF≈1 were represented by white circles, which indicate insufficient evidence for either hypotheses, and BF<1 were represented by grey circles and indicate evidence for null hypothesis (i.e., zero-correlation). Topographies above each panel depict Bayes Factors associated with a spatial searchlight analysis for each model’s partial correlation. For all models, the strongest contributions are evident over posterior electrodes. This suggests that eye-movements (reflected in frontal channel activity) are unlikely to have made a substantial contribution to the model performances.

For the word-based object dimension models, we found evidence for a partial correlation with the Perceptual-word model and Experiment 1 EEG RDM starting from 124 ms, which coincides with its peak. The Conceptual-word model also correlated with this EEG dataset from 176 ms, with a distinct single peak occurring at 196 ms. From this, we inferred that the Conceptual-word model had a later correlation onset and peak compared to the Perceptual-word model, mirroring the pattern observed in the image models. Furthermore, compared to the image models, the word models generally had a later onset and peak of partial correlation (see Table 1 for correlation onsets and peaks for each model). Finally, there was either insufficient evidence or evidence of no partial correlation between the Contextual-word model and the Experiment 1 EEG RDM.

**Table 1.** Partial Correlation Onsets and Peaks. The time points (ms) corresponding to correlation onsets (top row) and peaks (bottom row) were identified using Bayes Factors (BFs) cut-offs (see Methods for details). N.E. denotes no evidence.

| Experiment 1 |  |  |  |  |  | Experiment 2 |  |  |  |  |  |
| --- | --- | --- | --- | --- | --- | --- | --- | --- | --- | --- | --- |
| Image-task |  |  | Word-task |  |  | Image-task |  |  | Word-task |  |  |
| Per-cept | Con-cept | Con-text | Per-cept | Con-cept | Con-text | Per-cept | Con-cept | Con-text | Per-cept | Con-cept | Con-text |
| 108 ms | 184 ms | N.E. | 124 ms | 176 ms | N.E. | 108 ms | 160 ms | 384 ms | 120 ms | 176 ms | N.E. |
| 192 ms | 188 ms | N.E. | 124 ms | 196 ms | N.E. | 204 ms | 220 ms | 732 ms | 368 ms | 212 ms | N.E. |

### 3.2 Correlating the behavioural models with the Experiment 2 neural

#### RDM

The right column of Figure 3 shows the partial correlations between the Experiment 2 EEG dataset (acquired at a 3.33 Hz image presentation rate) and the behavioural models derived from the image-based triplet similarity task (Figure 3C) and the word-based triplet similarity task (Figure 3D). For the image-based models, we found evidence of a partial correlation between this neural RDM and the Perceptual-image mode beginning 108 ms after stimulus onset and peaking at 204 ms. The Conceptual-image model displayed a correlation onset of 160 ms, reaching its peak at 220 ms. Notably, we also found modest evidence of a partial correlation between the Contextual-image model and Experiment 2 neural RDM, starting from 384 ms and peaking at 732 ms. These results show that the correlation onset of the Conceptual-image model occurred relatively later compared to the Perceptual-image model, but the peak correlation occurred around a similar time for both models. In contrast, the modest unique contribution of the Contextual-image model arose rather later during the neural response.

Figure 3D shows the partial correlations between Experiment 2 neural RDM and the three models derived from word-based similarity judgements. The Perceptual-word model’s partial correlation started from 120 ms and peaked at 368 ms, while the Conceptual-word model correlated with EEG from 176 ms with a single distinct peak at 212 ms. For the Contextual-word model, we found either evidence for no correlation, or else insufficient evidence to support either outcome.

### 3.3. Summary of EEG Datasets

In summary, the object dimension models that uniquely explained the dynamic neural representation of the stimuli were largely consistent across both EEG datasets. Specifically, the perceptual and conceptual models, whether derived from image-based or word-based similarity judgements, consistently accounted for the neural response to object images in both Experiment 1 and 2. Evidence for a unique contribution from the contextual models was comparatively much weaker – indeed, the only unique contribution we observed was from the Contextual-image model to the Experiment 2 neural dataset. In this case, explanatory power both arose and peaked at relatively at late time points (384 ms and 732 ms respectively). All other statistical analyses provided insufficient evidence or evidence of no partial correlation between the contextual models and EEG.

From these results, we concluded that the pattern observed in Experiment 2 reflected a ‘scaled-up’ version of that seen for Experiment 1, with similar timings of correlation onsets and peak time points in the perceptual and conceptual models. This is likely due to the threefold increase in image repeats in Experiment 2, which enable a higher signal-to-noise ratio than the single image presentations that characterised Experiment 1. Given the high consistency of findings across the two EEG datasets, and considering the enhanced power provided by Experiment 2, our subsequent analyses focused exclusively on the neural data acquired in Experiment 2. Within this dataset, we pursued a series of more focused analysis questions detailed below.

### 3.4 Perceptual versus Conceptual models in explaining neural responses

In Figure 4A, we examined the relative contributions of the perceptual and conceptual models to explaining the neural representational structure of the neural RDM from Experiment 2 by subtracting the partial correlation of the conceptual model from the perceptual model, separately for each task modality. A two-tailed Bayesian *t*-test was used to compare the resulting difference profile to zero.

**Figure 4.**
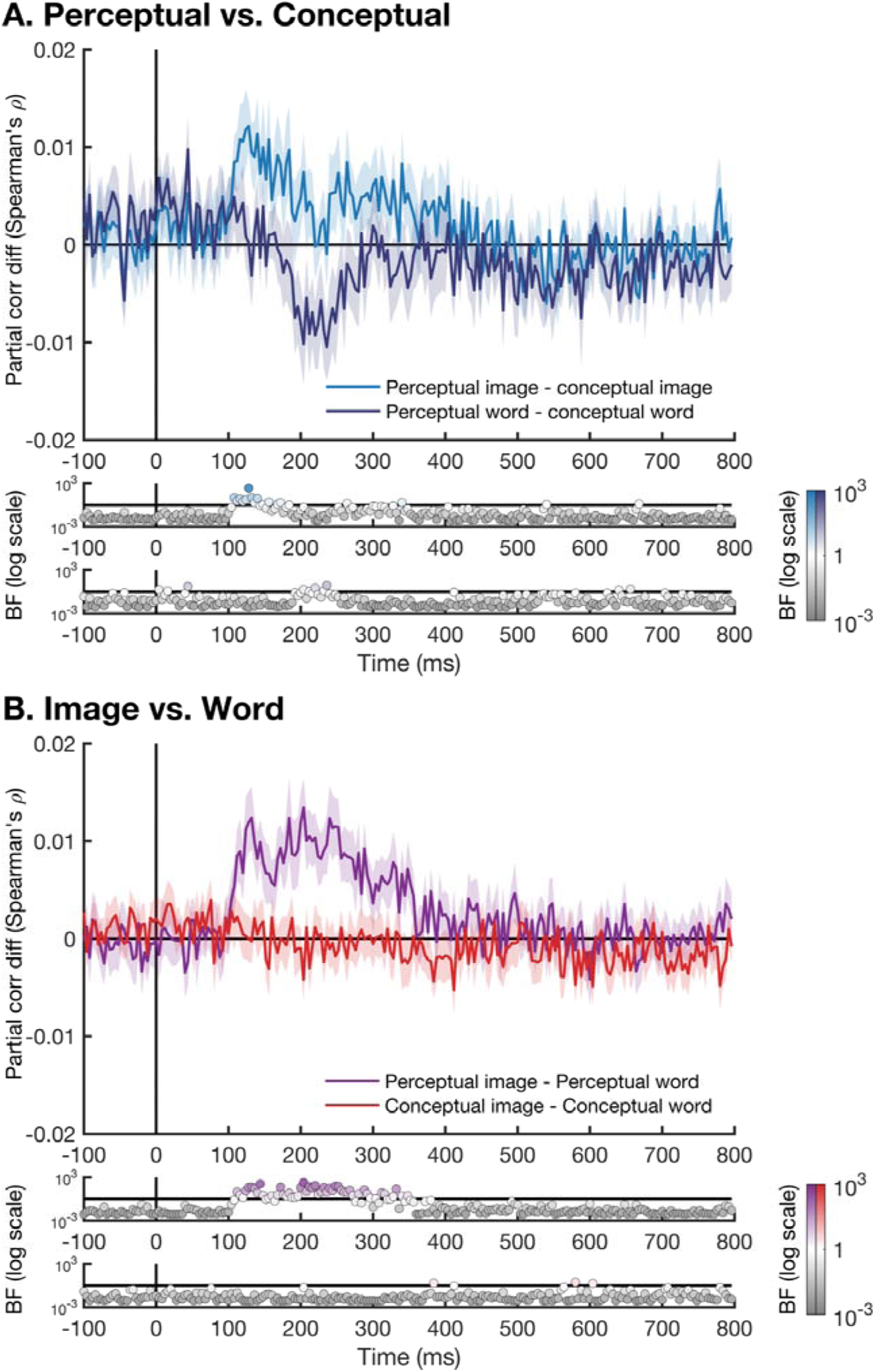
Spearman’s *ρ* differences contrasting the predictive power for various behavioural models. **(A)** Perceptual – Conceptual partial correlation differences calculated for each modality (i.e., Image or Word) separately. **(B)** Image – Word partial correlation differences calculated for each similarity dimension (i.e., Perceptual or Conceptual) separately. Shaded areas indicate standard error. Lower panels are log-scaled Bayes Factors testing difference scores ≠ 0.

When we compared the Perceptual-image model with the Conceptual-image model, their difference score exceeded zero during an early time window (108-140 ms), indicating that the Perceptual-image model provided a better explanation of the neural response to object images than the Conceptual-image model during this early phase. Outside this window, there was either no evidence or insufficient evidence of a difference between the two models, suggesting that both models performed equally well at explaining EEG during this period.

Comparing the performance of the Perceptual-word with the Conceptual-word model, only a few isolated time points showed evidence of a difference. Since no two or more consecutive time points showed such evidence, we concluded that there was limited evidence of difference between the Perceptual-word and Conceptual-word models, indicating that both models provided comparably good explanations of the EEG data across all time points.

In Figure 4B, we compared the modality in which the triplet similarity task was presented (i.e., image vs. word) within the same object dimension (i.e., perceptual or conceptual). When comparing the two perceptual models, we observed that the Perceptual-image model had a greater unique explanation of EEG neural response compared to the Perceptual-word model, during the period of 112 to 332 ms, although included a few time points that showed insufficient evidence. In contrast, there was no sustained evidence of a difference between the Conceptual-image and Conceptual-word models. Note that while the image-based and word-based conceptual models appeared to explain the neural data equally well, this need not imply that these two models captured the *same* conceptual information about objects (for further details on this, see Figure S1 in Supplementary Materials).

Overall, our analysis of the difference between the model performances suggests that: (1) the Perceptual-image model outperformed the Conceptual-image model at early time points; (2) the Perceptual-image model was better predictor of the (image-derived) EEG responses compared to the Perceptual-word model across a broad time window; (3) there was limited evidence of a difference in explanatory power between the Perceptual-word and Conceptual-word models, as well as the Conceptual-image and Conceptual-word models.

### 3.5 Do Contextual models play any role in explaining the neural response to objects?

In a final analysis, we focused specifically on the models of objects’ contextual attributes, which thus far show comparatively weaker capacity to predict the neural response to object images (Contextual-image model correlating with the Experiment 2 neural RDM at isolated late time points; see Figure 3C and Table 1). At this stage, it remains unclear whether behaviourally derived models of objects’ contextual associations lack explanatory power altogether, or if their explanatory power is simply not unique, due to shared variance with the perceptual and/or conceptual models. To examine this, we calculated the partial correlations between each contextual model and the Experiment 2 neural RDM while partialling out the perceptual and conceptual models in separate analyses. As a baseline, we also calculated the full correlation between each contextual model and the neural RDM (i.e., without removing shared variance with any other model). In doing so, we aimed to determine whether the contextual model considered in isolation could account for any variance in the EEG data, and if so, clarify the extent to which this variance overlapped with that explained by the perceptual and conceptual models individually.

Figure 5A shows the full and partial correlations of the Contextual-image model with EEG. We found evidence that the Contextual-image model did correlate with the EEG data when no other models were taken into account (onset at 120 ms and peaking at 244 ms). When the variance explained by the Perceptual-image model was partialled out, we retained evidence of a partial correlation between the Contextual-image model and EEG from 140 ms onwards. When the covariate was changed to be the Conceptual-image model, we retained evidence of a partial correlation between the Contextual-image model and EEG during the late time window from 384 ms. This would suggest that the Conceptual-image model shared a similar amount of variance with the Contextual-image model, and that when variance accounted for by each were taken into consideration, the Contextual-image model retained little unique explanatory power.

**Figure 5.**
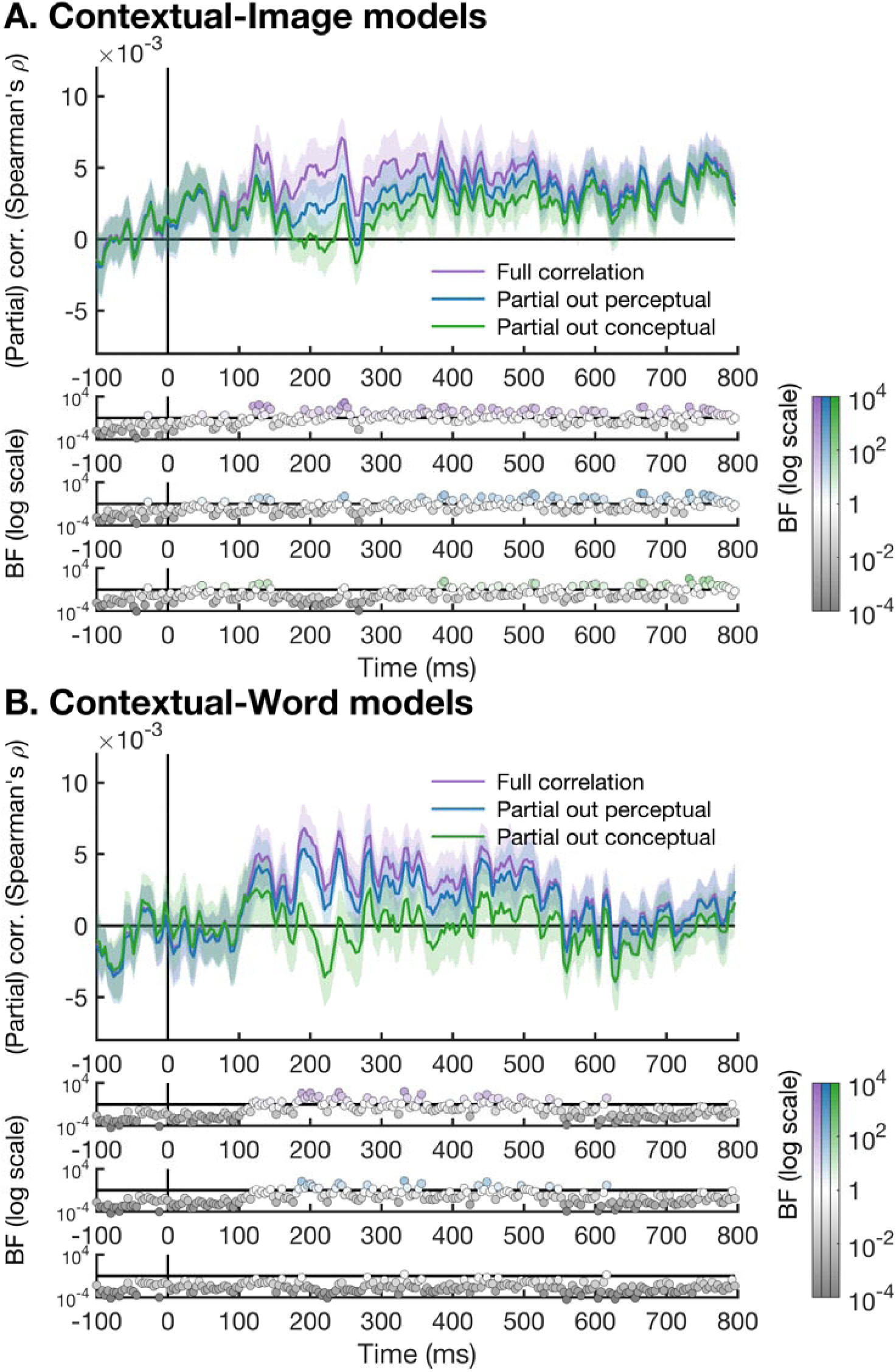
Full and Partial Correlations for the Contextual Models based on **(A)** Image-task and **(B)** Word task. The lines were temporally smoothed with a kernel of 3 samples (12 ms) for visibility. Below each plot, Bayes factors (BFs) appear for the full correlation (top), partial correlation controlling for the perceptual model within the same modality (middle) and partial correlation controlling for the conceptual model within the same modality (bottom). The Bayes factors (BFs) were based on unsmoothed data.

Figure 5B shows the full and partial correlations of the Contextual-word model with EEG. Evidence indicated that the Contextual-word model began correlating with EEG at 184 ms. When the Perceptual-word model was included as a covariate and partialled out from the full correlation, the correlations were mostly preserved, with a shifted onset of 196 ms. This suggested minimal shared variance between the Contextual-word and the Perceptual-word models. However, when the covariate was switched to the Conceptual-word model, there was no longer any evidence supporting the Contextual-word model’s unique explanatory power, in contrast to the results when the Perceptual-word model was partialled out. In general, these findings suggest that the contextual models shared relatively more variance with the conceptual models than the perceptual models, particularly in the case of word-based models. While the contextual models do explain variance in the EEG data, they do not provide unique explanatory power beyond what is captured by the conceptual models.

## 4. DISCUSSION

This study investigated how contextual, perceptual, and conceptual object attributes contribute to the neural responses evoked by visual objects. We used representational similarity analysis (RSA) to correlate two EEG datasets recorded during the passive viewing of object images with behavioural models based on similarity judgements of objects’ perceptual, conceptual, or contextual properties. Separate behavioural models were derived for image and word label versions of the objects. Our findings revealed that behavioural similarity judgements along the perceptual and conceptual dimensions accounted for independent aspects of the neural response to object images. Our results underscore the robustness of object representations in the human brain, as the temporal dynamics of perceptual and conceptual object feature models’ explanatory power were consistent across both word-based and image-based models, and across EEG datasets with differing signal-to-noise ratios due to differences in presentation rate, stimulus repetition, and inter-stimulus interval. Notably, there was no evidence that contextual object attributes captured by in our behavioural models were meaningfully related to the neural response elicited by object images over and above what could be explained by the perceptual and conceptual models.

### A perceptual-conceptual continuum of object features across both image and word modalities

A key finding from our study is that the perceptual features of objects deemed relevant by human observers dominated the early stages of the neural response to object images, with conceptual aspects of objects better explaining a later stage processing. Information about objects’ perceptual attributes emerged as early as approximately 100 ms after stimulus onset, while conceptual information was reflected in the neural response to object images from approximately 160 ms onwards. This pattern was consistent regardless of the image presentation rate, and evident using models derived from both the image and word modalities, although onsets were slightly later for the Perceptual-word compared to image models (see Table 1). These findings largely align with established object recognition literature that has identified early processing of lower level visual features (such as object shape and colour) followed by abstract conceptual attributes (such as object category) (Carlson et al., 2013; Cichy et al., 2014; Contini et al., 2017, 2020; Grootswagers et al., 2019a, 2019b; Teichmann et al., 2026).

We found that perceptual similarity between object images uniquely explained EEG neural response from approximately 100 ms after stimulus onset, peaking at approximately 200 ms for both Experiment 1 and 2. This is in line with previous studies that have compared behavioural models of perceived visual similarity to EEG responses to visual stimuli (Cichy et al., 2019; Wardle et al., 2016). Cichy and colleagues found the unique emergence of visual features such as colour and shape at 82 ms and 183 ms respectively, peaking at 82-117 ms and 183-202 ms. Similarly, Wardle et al. found that the perceived similarity of artificial shapes explained neural response from as early as 50 ms, with the highest correlation observed at 150 ms.

Although variations in experimental designs and stimuli make direct comparisons of correlation onsets and peaks between studies challenging (Contini et al., 2017), our findings are consistent with the broader agreement that visual features underlying behaviour are processed within the first 200 ms of visual object recognition (Carlson et al., 2013; Cichy et al., 2014; Contini et al., 2017). While we cannot determine which specific visual features participants focused on when evaluating objects’ perceptual similarity, the Perceptual-image model demonstrated the earliest time point of unique correlation among all behavioural models, which suggest that participants likely relied on lower-level properties of objects such as colour or shape to guide their similarity judgements. The temporal dynamics associated with conceptual attributes of object images observed here also concord with extant findings. In our case, the Conceptual-image model uniquely explained the EEG response from approximately 160 ms onwards, peaking around 200 ms, a relatively later onset but similar peak compared to the Perceptual-image model. The partly overlapping time points of correlation between the perceptual and conceptual image models highlight the dynamic evolution of representations in the brain, transitioning from lower-level perceptual to higher-level semantic properties over the first 200 ms after stimulus onset (Bankson et al., 2018; Clarke et al., 2013, 2015).

The inclusion of word-based models of object dimensions enabled us to inspect the relative contribution of perceptual, conceptual, and contextual object properties to the neural response while controlling for category-nonspecific visual features of individual stimulus images. Here we found that the Perceptual-word model held unique explanatory power for our neural data from approximately 120 ms after stimulus onset. This suggests that perceptual object properties accessible via object labels alone are represented during the early stages of visual object perception, a finding that may be underpinned by mental imagery. Specifically, when participants in our behavioural perceptual similarity task read a word like "AIRPLANE", they may well have activated this concepts’ associated visual features (e.g., shape, colour) in their minds to help guide their judgement of this object-concept’s similarity with the other concepts on that trial (Pearson et al., 2015). Research on mental imagery indicates that imagining a visual stimulus activates the visual cortex, engaging similar neural mechanisms to those involved in perceiving an actual object (Kosslyn et al., 1995; Mechelli et al., 2004; O’Craven & Kanwisher, 2000). At the same time, vivid visual imagery is certainly not required to do the word-based perceptual similarity task – judgements in this case could also be based on internally constructed representations of the object-labels. Regardless of underlying processes that behavioural participants used, it remains the case that visual processing of objects as early as 120 ms following stimulus onset could be explained by the perceptual properties that online participants used when considering the similarity of object-labels. Consistent with this is the finding that the word-based Perceptual model’s correlation onset and peak arose later than those for the image-based Perceptual model, which suggests that the specific perceptual properties tapped by object-label similarity judgements are somewhat more abstract than those tapped by image-based judgements.

The relatively later onset of the Conceptual-word model’s explanation of EEG (approximately 180 ms after stimulus onset) further suggests that the specific abstract/semantic features of objects that observers access via object labels are represented in the neural response to objects somewhat later than their stereotypically-associated visual features. Taken together, these results suggest that reading object-labels can activate both perceptual and conceptual object knowledge, but in a different manner to when observers encounter an object image, extracting visual features from the stimuli and derive their conceptual meaning.

A unique aspect of our study is its direct comparison of behaviourally-derived object dimension models based on judgements elicited by word and image versions of the same object concepts. Our findings suggest there is significant overlap in the object knowledge accessed via object image and word labels, in that both contribute to explaining the neural activity involved in visual object recognition. However, there appears to be a trend where lexically-accessed visual and conceptual features explain neural processes at later time points compared to image-derived object knowledge for the perceptual models. In demonstrating that lexically-accessed perceptual and conceptual features can also explain the neural responses to visual objects, we add to an existing literature that has mostly relied on object images to generate behavioural object dimension models (Cichy et al., 2019; Giari et al., 2020). The findings here are some of the first to capture how presentation modality (word vs. image) differentially activates object knowledge within the same dimension, with object labels facilitating access to semantic aspects of object that are less visually confounded than those elicited by object images (Giari et al., 2020). Precedent for our approach does exist in terms of neural responses, with Giari and colleagues. measuring neural responses for image and word versions of the same objects with EEG. Specifically, they compared conceptual object similarity with the representational structure of MEG responses elicited by both pictures and words, to delineate the time course of higher-level semantic processing evoked by reading object labels (which are not subject to the visual-conceptual covariance present in object images). Several factors distinguish our study from that of Giari et al.: where they used a category-typicality rating task during MEG recording to activate conceptual object representations, the neural representations of objects in our data were acquired under passive viewing conditions. That is, we deliberately did not direct attention to object features, but rather engaged participants in an orthogonal fixation colour change detection task that rendered the objects task-irrelevant. That we find evidence of conceptual processing using both an orthogonal task and a faster presentation rate than Giari et al. (100 ms vs. 400 ms) highlights the rapid and automatic nature of object concept activations during visual perception.

### Interpretation considerations

While the partial correlation values we report here are relatively small in absolute magnitude, within a multivariate decoding framework, decoding accuracy and correlation strength are understood to depend strongly on multivariate analysis choices (e.g., trial averaging, cross-validation scheme, and representational format), and therefore do not map straightforwardly onto classical effect size measures such as Cohen’s d (e.g., Allefeld & Haynes, 2014; Combrisson & Jerbi, 2015; Dubois et al., 2015; Hebart & Baker, 2018). Indeed, even accuracies or correlations close to chance can carry meaningful information if they are reliable and generalise across participants and timepoints (Christophel et al., 2015). In the present case, our purpose was to assess whether various behavioural models explained variance in time-resolved EEG representational structure i) above chance and ii) relative to other models. In this context, small but reliable correlations are expected, particularly given the noisy nature of single-trial EEG data and the fact that we correlate neural RDMs against high-level behavioural models rather than stimulus labels. It is compelling to find that even in this context, the evidence supporting the unique predictive power of our models is very strong, with Bayes Factors exceeding 100,000 in some cases (see Figure 3).

Additionally, it is also relevant to consider what role eye movements could have played in explaining our results (Linde-Domingo & Spitzer, 2024; Mostert et al., 2018; Quax et al., 2019). Happily, the current design limits participants’ opportunity to make meaningful saccades via a combination of very short presentation durations, parafoveal stimulus presentation, and a focal orthogonal task. Nevertheless, we cannot completely exclude the possibility that there could be a systematic relationship between participants’ eye movements or blinks and the perceptual, conceptual, or contextual properties of our visual stimuli. Yet the results of our channel-based searchlight analysis would argue against this possibility: the dominant contribution to all our models arises at posterior sensors (where we expect visual neural responses to be strongest), rather than frontal electrodes that might hint at eye-movement-related activity.

### Contrasting perceptual and conceptual models within each modality

When comparing the performance of different models based on object dimension or modality against the EEG neural data, we found that the Perceptual-image model outperformed the Conceptual-image model at early time points (approximately 100-140 ms after stimulus onset). The early dominance of the Perceptual-image model over the Conceptual-image model further supports our findings from the individual partial correlations of the behavioural models across the two modalities and EEG datasets, indicating that perceptual features are processed earlier than conceptual features. There was limited evidence of a difference in explanatory power between the Perceptual-word and Conceptual-word models. However, this still suggests a general pattern of later processing of conceptual features compared to perceptual features, consistent with the Perceptual-image and Conceptual-image models. Taken together, this suggests that for image-derived behavioural models, perceptual features explain EEG data better than conceptual features. The same was not true for the word-based models, where we observed a trend for the Conceptual-word model explaining the neural responses better than the Perceptual-word model, although not reaching the threshold for evidence. Future studies could explore the factors contributing to this modality-specific distinction, particularly in how perceptual and conceptual features are evoked differently by pictorial and lexical stimuli, which could offer more insights into the temporal dynamics of perceptual and conceptual feature integration in the brain.

### Contrasting image- and word-based models within each object dimension

The superior fit of the Perceptual-image model compared to the Perceptual-word model across a broad time range of approximately 100-330 ms suggests that perceptual features evoked by directly viewing object images better explain the EEG than those evoked by reading word labels. As discussed earlier, reading word labels requires a degree of mentalisation to evoke perceptual features, which may vary significantly across participants. The variability likely limits the explanatory power of the Perceptual-word model compared to the Perceptual-image model. In contrast, there was no evidence that the two conceptual models based on different modalities differed in their explanatory power. Moreover, incorporating the Conceptual-word and Conceptual-image models into a single analysis did not reduce either model’s partial correlation to zero (Figure S1 in Supplementary Materials), raising the possibility that the two captured somewhat distinct elements of objects’ conceptual attributes. This could reflect different ways of accessing conceptual knowledge: image-cued activation may involve processing visual features to retrieve associated conceptual attributes, while word-cued activation relies on lexical processing to access the conceptual knowledge of the object stored in the brain.

### Contextual object associations do not uniquely predict neural responses to objects

In contrast to the relatively clear connection between the perceptual and conceptual models and the neural data, we found only fleeting evidence that the models of objects’ contextual associations uniquely explained the neural response to object images. This evidence was limited to Experiment 2 neural dataset, and corresponded to timepoints from 384 ms onwards. The minimal explanatory power of the contextual models was surprising given that prior studies have found that observers are highly sensitive to violations of object context (Bar, 2004; Biederman et al., 1973; Davenport & Potter, 2004; Ganis & Kutas, 2003; Mudrik et al., 2010; Mudrik et al., 2014; Oliva & Torralba, 2007; Quek & Peelen, 2020; Võ et al., 2019), and recognise objects more efficiently when they appear alongside related objects (Auckland et al., 2007; Bar & Ullman, 1996; Davenport, 2007; Green & Hummel, 2006; Kaiser et al., 2014, 2019; Quek et al., 2025). As such, we might well have expected information about objects’ contextual associations might well be carried in the evoked responses to object images.

One possible explanation for the lack of unique explanatory power for our contextual attribute models concerns the overlap in conceptual and contextual features of our selected 190 object concepts. Indeed, although both the Contextual-word and Contextual-image model *did* predict neural responses when considered in isolation (i.e., without other models as covariates), they lost most of their explanatory power when the conceptual model from the same modality was included as a covariate. Previous studies have found that the parahippocampal cortex encodes the contextual representation of objects, specifically how certain objects tend to co-occur with one another within a context (Aminoff et al., 2013; Bar, 2004). Martin and colleagues also found in their fMRI-RSA analysis that conceptual representations were activated in parahippocampal cortex when participants were engaged in a task that evoked conceptual features (Martin et al., 2018). In line with their findings, our results also seem to demonstrate that objects that tend to occur in the same context often share many conceptual features but can be visually distinct (e.g., a toothbrush and toothpaste). Our study also extends this further by demonstrating that in time-resolved whole-brain EEG response to object images, contextual features, whether derived from images or words, were not uniquely processed by the brain on top of conceptual features; rather, they were processed as part of conceptual processing. Another, non-mutually-exclusive possibility is that contextual associations between object concepts are represented most strongly when two (or more) objects are encountered simultaneously (as has often been the case in prior studies examining contextual relations between objects). Future studies seeking to disentangle the neural timecourse of contextual object knowledge from its perceptual and conceptual counterparts might exploit such multi-object processing paradigms (Quek et al., 2025; Quek & Peelen, 2020), in which the strength of conceptual and contextual object associations could be strictly orthogonalised.

## Conclusion

Our study demonstrates the brain’s robust and cascaded integration of perceptual and conceptual features in forming object representations. Using RSA to correlate EEG responses with behavioural models, we found that perceptual features dominate early neural responses, while conceptual features emerge later. These findings were consistent across different behavioural task modalities and EEG datasets. Notably, although contextual associations between objects did meaningfully predict the structure of neural object representations, this explanatory power overlapped significantly with models of conceptual object features. By using functionally relevant models of different object dimensions, this work provides insight into how these dimensions are generally represented when observers look at objects. Together, these findings contribute to our understanding of the brain’s dynamic processing of visual objects.

## Acknowledgements

This project was partially funded by an Australian Research Council Discovery Project grant (DP200101787) awarded to T. A. Carlson

## Notes

https://osf.io/jy284/

### Competing Interest Statement

The authors have declared no competing interest.

### Summary of Updates

Manuscript updated with changes in-text. One analysis moved to supplemental materials. Scalp topographies added to the results.

## REFERENCES

Almeida, J., Fracasso, A., Kristensen, S., Valério, D., Bergström, F., Chakravarthi, R., … Walbrin, J. (2023). Neural and behavioral signatures of the multidimensionality of manipulable object processing. Communications Biology, 6(1), 940. doi:10.1038/s42003-023-05323-x

Aminoff, E. M., Kveraga, K., & Bar, M. (2013). The role of the parahippocampal cortex in cognition. Trends in Cognitive Sciences, 17(8), 379–390. doi:10.1016/j.tics.2013.06.009

Auckland, M. E., Cave, K. R., & Donnelly, N. (2007). Nontarget objects can influence perceptual processes during object recognition. Psychonomic Bulletin & Review, 14(2), 332–337.

Bankson, B. B., Hebart, M. N., Groen, I. I. A., & Baker, C. I. (2018). The temporal evolution of conceptual object representations revealed through models of behavior, semantics and deep neural networks. Neuroimage, 178, 172–182. 10.1016/j.neuroimage.2018.05.037

Bar, M. (2004). Visual objects in context. Nature Reviews Neuroscience, 5(8), 617.

Bar, M., & Ullman, S. (1996). Spatial Context in Recognition. Perception, 25(3), 343–352. doi:10.1068/p250343

Biederman, I., Glass, A. L., & Stacy, E. W. (1973). Searching for objects in real-world scenes. Journal of Experimental Psychology, 97(1), 22–27. doi:10.1037/h0033776

Brandman, T., & Peelen, M. V. (2017). Interaction between Scene and Object Processing Revealed by Human fMRI and MEG Decoding. Journal of Neuroscience, 37(32), 7700–7710. doi:10.1523/jneurosci.0582-17.2017

Carlson, T., Tovar, D. A., Alink, A., & Kriegeskorte, N. (2013). Representational dynamics of object vision: the first 1000 ms. Journal of Vision, 13(10), 1.

Cichy, R. M., Kriegeskorte, N., Jozwik, K. M., van den Bosch, J. J. F., & Charest, I. (2019). The spatiotemporal neural dynamics underlying perceived similarity for real-world objects. Neuroimage, 194, 12–24. 10.1016/j.neuroimage.2019.03.031

Cichy, R. M., Pantazis, D., & Oliva, A. (2014). Resolving human object recognition in space and time. Nature Neuroscience, 17(3), 455–462. doi:10.1038/nn.3635

Clarke, A., Taylor, K. I., Devereux, B., Randall, B., & Tyler, L. K. (2012). From Perception to Conception: How Meaningful Objects Are Processed over Time. Cerebral Cortex, 23(1), 187–197. doi:10.1093/cercor/bhs002

Clarke, A., & Tyler, L. K. (2015). Understanding What We See: How We Derive Meaning From Vision. Trends in Cognitive Sciences, 19(11), 677–687. doi:10.1016/j.tics.2015.08.008

Contini, E. W., Goddard, E., Grootswagers, T., Williams, M., & Carlson, T. (2020). A humanness dimension to visual object coding in the brain. Neuroimage, 117139. 10.1016/j.neuroimage.2020.117139

Contini, E. W., Wardle, S. G., & Carlson, T. A. (2017). Decoding the time-course of object recognition in the human brain: From visual features to categorical decisions. Neuropsychologia, 105, 165–176. 10.1016/j.neuropsychologia.2017.02.013

Davenport, J. L. (2007). Consistency effects between objects in scenes. Memory & Cognition, 35(3), 393–401. doi:10.3758/bf03193280

Davenport, J. L., & Potter, M. C. (2004). Scene Consistency in Object and Background Perception. Psychological Science, 15(8), 559–564. doi:10.1111/j.0956-7976.2004.00719.x

Delorme, A., & Makeig, S. (2004). EEGLAB: an open source toolbox for analysis of single-trial EEG dynamics including independent component analysis. Journal of Neuroscience Methods, 134(1), 9–21. 10.1016/j.jneumeth.2003.10.009

Edelman, S. (1998). Representation is representation of similarities. Behavioral and Brain Sciences, 21(4), 449–467. doi:10.1017/S0140525X98001253

Ganis, G., & Kutas, M. (2003). An electrophysiological study of scene effects on object identification. Brain Research: Cognitive Brain Research, 16(2), 123–144. doi:10.1016/s0926-6410(02)00244-6

Giari, G., Leonardelli, E., Tao, Y., Machado, M., & Fairhall, S. L. (2020). Spatiotemporal properties of the neural representation of conceptual content for words and pictures – an MEG study. Neuroimage, 219, 116913. 10.1016/j.neuroimage.2020.116913

Green, C., & Hummel, J. E. (2006). Familiar interacting object pairs are perceptually grouped. Journal of Experimental Psychology: Human Perception and Performance, 32(5), 1107.

Grootswagers, T., Robinson, A. K., & Carlson, T. (2019a). The representational dynamics of visual objects in rapid serial visual processing streams. Neuroimage, 188, 668–679. 10.1016/j.neuroimage.2018.12.046

Grootswagers, T., Robinson, A. K., Shatek, S. M., & Carlson, T. A. (2019b). Untangling featural and conceptual object representations. Neuroimage, 202, 116083.

Grootswagers, T., Wardle, S. G., & Carlson, T. (2017). Decoding dynamic brain patterns from evoked responses: A tutorial on multivariate pattern analysis applied to time series neuroimaging data. Journal of Cognitive Neuroscience, 29(4), 677–697. doi:10.1162/jocn_a_01068

Grootswagers, T., Zhou, I., Robinson, A. K., Hebart, M. N., & Carlson, T. A. (2022). Human EEG recordings for 1,854 concepts presented in rapid serial visual presentation streams. Scientific Data, 9(1), 3. doi:10.1038/s41597-021-01102-7

Hebart, M. N., Contier, O., Teichmann, L., Rockter, A. H., Zheng, C. Y., Kidder, A., … Baker, C. I. (2023). THINGS-data, a multimodal collection of large-scale datasets for investigating object representations in human brain and behavior. eLife, 12, e82580. doi:10.7554/eLife.82580

Hebart, M. N., Dickter, A. H., Kidder, A., Kwok, W. Y., Corriveau, A., Van Wicklin, C., & Baker, C. I. (2019). THINGS: A database of 1,854 object concepts and more than 26,000 naturalistic object images. PloS One, 14(10), e0223792. doi:10.1371/journal.pone.0223792

Hebart, M. N., Zheng, C. Y., Pereira, F., & Baker, C. I. (2020). Revealing the multidimensional mental representations of natural objects underlying human similarity judgements. Nature Human Behaviour, 4(11), 1173–1185. doi:10.1038/s41562-020-00951-3

Isik, L., Meyers, E. M., Leibo, J. Z., & Poggio, T. (2014). The dynamics of invariant object recognition in the human visual system. Journal of Neurophysiology, 111(1), 91–102. doi:10.1152/jn.00394.2013

Kaiser, D., Quek, G. L., Cichy, R. M., & Peelen, M. V. (2019). Object Vision in a Structured World. Trends in Cognitive Sciences, 23(8), 672–685. doi:10.1016/j.tics.2019.04.013

Kaiser, D., Stein, T., & Peelen, M. V. (2014). Object grouping based on real-world regularities facilitates perception by reducing competitive interactions in visual cortex. Proceedings of the National Academy of Sciences of the United States of America, 111(30), 11217–11222.

Kok, P., Brouwer, G. J., van Gerven, M. A. J., & de Lange, F. P. (2013). Prior Expectations Bias Sensory Representations in Visual Cortex. The Journal of Neuroscience, 33(41), 16275–16284. doi:10.1523/jneurosci.0742-13.2013

Kosslyn, S. M., Thompson, W. L., Klm, I. J., & Alpert, N. M. (1995). Topographical representations of mental images in primary visual cortex. Nature, 378(6556), 496–498. doi:10.1038/378496a0

Kriegeskorte, N., Mur, M., & Bandettini, P. (2008). Representational Similarity Analysis – Connecting the Branches of Systems Neuroscience. Frontiers in Systems Neuroscience, 2, 4. doi:10.3389/neuro.06.004.2008

Martin, C. B., Douglas, D., Newsome, R. N., Man, L. L. Y., & Barense, M. D. (2018). Integrative and distinctive coding of visual and conceptual object features in the ventral visual stream. eLife, 7, e31873. doi:10.7554/eLife.31873

Mechelli, A., Price, C. J., Friston, K. J., & Ishai, A. (2004). Where Bottom-up Meets Top-down: Neuronal Interactions during Perception and Imagery. Cerebral Cortex, 14(11), 1256–1265. doi:10.1093/cercor/bhh087

Moerel, D., & Grootswagers, T. (2025). Reconstruction of Partial Dissimilarity Matrices for Cognitive Neuroscience. arXiv preprint arXiv:2506.00484.

Moerel, D., Psihoyos, J., & Carlson, T. A. (2024). The Time-Course of Food Representation in the Human Brain. The Journal of Neuroscience, 44(26), e1101232024. doi:10.1523/jneurosci.1101-23.2024

Morey, R. D., & Rouder, J. N. (2011). Bayes factor approaches for testing interval null hypotheses. Psychological Methods, 16(4), 406–419. doi:10.1037/a0024377

Morey, R. D., Rouder, J. N., Jamil, T., & Morey, M. R. D. (2015). Package ‘bayesfactor’. In.

Mudrik, L., Lamy, D., & Deouell, L. Y. (2010). ERP evidence for context congruity effects during simultaneous object–scene processing. Neuropsychologia, 48(2), 507–517.

Mudrik, L., Shalgi, S., Lamy, D., & Deouell, L. Y. (2014). Synchronous contextual irregularities affect early scene processing: Replication and extension. Neuropsychologia, 56, 447–458. 10.1016/j.neuropsychologia.2014.02.020

Nosofsky, R. M. (1986). Attention, similarity, and the identification-categorization relationship. Journal of Experimental Psychology: General, 115(1), 39–61. doi:10.1037//0096-3445.115.1.39

O’Craven, K. M., & Kanwisher, N. (2000). Mental Imagery of Faces and Places Activates Corresponding Stimulus-Specific Brain Regions. Journal of Cognitive Neuroscience, 12(6), 1013–1023. doi:10.1162/08989290051137549

Oliva, A., & Torralba, A. (2007). The role of context in object recognition. Trends in Cognitive Sciences, 11(12), 520–527. 10.1016/j.tics.2007.09.009

Oostenveld, R., & Praamstra, P. (2001). The five percent electrode system for high-resolution EEG and ERP measurements. Clinical Neurophysiology, 112(4), 713–719. 10.1016/S1388-2457(00)00527-7

Oosterhof, N. N., Connolly, A. C., & Haxby, J. V. (2016). CoSMoMVPA: Multi-Modal Multivariate Pattern Analysis of Neuroimaging Data in Matlab/GNU Octave. Frontiers in Neuroinformatics, 10. doi:10.3389/fninf.2016.00027

Pearson, J., Naselaris, T., Holmes, E. A., & Kosslyn, S. M. (2015). Mental Imagery: Functional Mechanisms and Clinical Applications. Trends in Cognitive Sciences, 19(10), 590–602. doi:10.1016/j.tics.2015.08.003

Peirce, J., Gray, J. R., Simpson, S., MacAskill, M., Höchenberger, R., Sogo, H., … Lindeløv, J. K. (2019). PsychoPy2: Experiments in behavior made easy. Behavior Research Methods, 51(1), 195–203. doi:10.3758/s13428-018-01193-y

Proklova, D., Kaiser, D., & Peelen, M. V. (2019). MEG sensor patterns reflect perceptual but not categorical similarity of animate and inanimate objects. Neuroimage, 193, 167–177. 10.1016/j.neuroimage.2019.03.028

Quek, G. L., & Peelen, M. V. (2020). Contextual and Spatial Associations Between Objects Interactively Modulate Visual Processing. Cerebral Cortex, 30(12), 6391–6404. doi:10.1093/cercor/bhaa197

Quek, G. L., Theodorou, A., & Peelen, M. V. (in press). The timecourse of inter-object contextual facilitation. Cortex. doi:10.1101/2023.05.30.542965

Teichmann, L., Hebart, M. N., & Baker, C. I. (2025). Dynamic representation of multidimensional object properties in the human brain. bioRxiv. doi:10.1101/2023.09.08.556679

Teichmann, L., Moerel, D., Baker, C., & Grootswagers, T. (2022). An Empirically Driven Guide on Using Bayes Factors for M/EEG Decoding. Aperture Neuro. doi:10.52294/ApertureNeuro.2022.2.MAOC6465

Võ, M. L. H., Boettcher, S. E. P., & Draschkow, D. (2019). Reading scenes: how scene grammar guides attention and aids perception in real-world environments. Current Opinion in Psychology, 29, 205–210. 10.1016/j.copsyc.2019.03.009

Wardle, S. G., Kriegeskorte, N., Grootswagers, T., Khaligh-Razavi, S.-M., & Carlson, T. A. (2016). Perceptual similarity of visual patterns predicts dynamic neural activation patterns measured with MEG. Neuroimage, 132, 59–70. 10.1016/j.neuroimage.2016.02.019

Wischnewski, M., & Peelen, M. V. (2021). Causal neural mechanisms of context-based object recognition. eLife, 10, e69736. doi:10.7554/eLife.69736

