## Supplemental Materials for "Disentangling objects’ contextual associations from perceptual and conceptual attributes using time-resolved neural decoding"

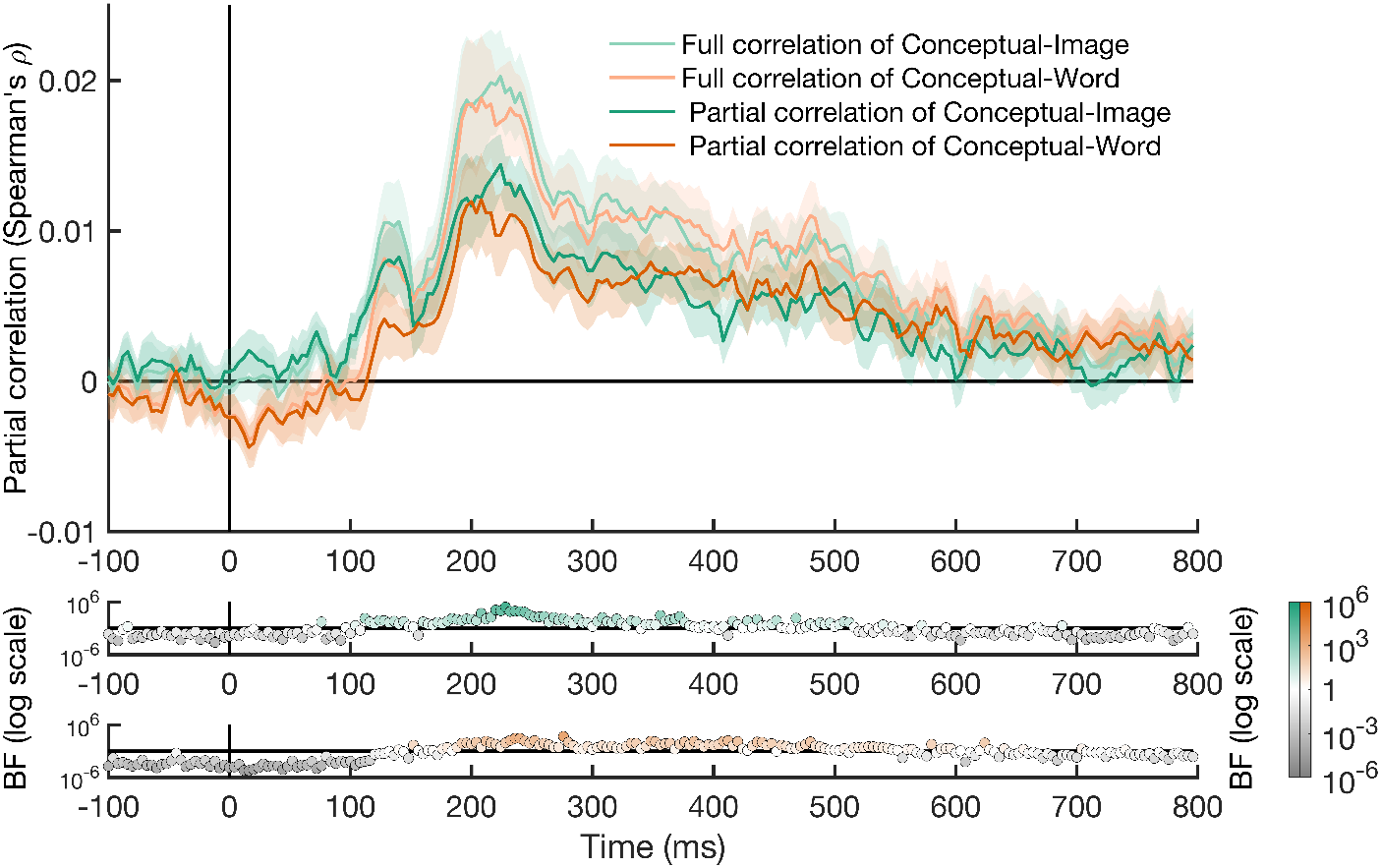

**Figure S1. Comparing Full and Partial Correlations of Conceptual-image and Conceptual-word models.** To consider whether the image- and word-based Conceptual models captured overlapping representational structure, we inspected both the full correlations between each conceptual model and the Experiment 2 neural RDM (pale lines) and the correlations remaining after each model was partialled out of the other (darker liners). While partial correlations for both models were reduced relative to full correlations, we found that removing the shared variance between the Conceptual-image and Conceptual-word model did not fully eliminate either model’s unique explanatory power. This suggests that the two conceptual models each had unique contributions for explaining the neural responses to objects, potentially capturing different aspects of conceptual object knowledge. This possibility would seem further underscored by the fact that although the Conceptual-image and Conceptual-word model were significantly correlated, they were by no means identical (see Figure 2). Correlations have been temporally smoothed with a kernel of 3 samples (12 ms) for visibility; Bayes factors (BFs) were based on unsmoothed data and are plotted only for the partial correlations.
